# SWI/SNF chromatin remodeling controls Notch-responsive enhancer accessibility

**DOI:** 10.1101/399501

**Authors:** Zoe Pillidge, Sarah J Bray

## Abstract

Notch signaling plays a key role in many cell fate decisions during development by directing different gene expression programs via the transcription factor CSL, known as Su(H) in *Drosophila.* Which target genes are responsive to Notch signaling is influenced by the chromatin state of enhancers, yet how this is regulated is not fully known. Detecting an increase in the histone variant H3.3 in response to Notch signaling, we tested which chromatin remodelers or histone chaperones were required for the changes in enhancer accessibility to Su(H) binding. This revealed a crucial role for the Brahma SWI/SNF chromatin remodeling complex in conferring enhancer accessibility and enabling the transcriptional response. The Notch-responsive regions had high levels of nucleosome turnover which were dependent on the Brahma complex, increased with Notch signaling and primarily involved histone H3.3. Together these results highlight the importance of SWI/SNF-mediated nucleosome turnover in rendering enhancers responsive to Notch.

## Introduction

Many cell fate decisions during development are directed by Notch signaling between neighboring cells, and misregulation of the pathway results in a variety of complex diseases [1,2]. Notch, the receptor, becomes cleaved upon binding to cell-surface ligands, freeing the Notch intracellular domain (NICD) to travel directly to the nucleus and activate target gene expression. Depending on the context, different genes are targeted by the Notch transcription complex due to differential enhancer accessibility [3–5]. Furthermore, successful activation involves largescale changes in histone modifications and chromatin accessibility across the target enhancers [4,6–8]. How these changes in chromatin structure are brought about remains to be determined.

The conserved DNA binding partner of NICD is known as Suppressor of Hairless (Su(H)) in *Drosophila melanogaster* or CSL more generally. In the absence of Notch activity, Su(H) partners with co-repressive proteins to prevent transcription, often acting on the same genes which are induced upon Notch signal activation [5]. The switch from genes being repressed (Notch-OFF) to activated (Notch-ON) involves a change in the dynamics of Su(H) binding, so that it acquires a longer residence time when participating in the activating complex, as well as increased accessibility of the DNA [8]. Several histone acetyltransferases and methyltransferases contribute to this switch [8–10] and their actions could explain some of the changes in histone post-translational modifications that have been observed [4,6,7]. However, the histone modifiers that have been identified do not explain how target enhancer accessibility is regulated, making it likely that other factors contribute.

One way that chromatin structure can be altered is by a change in the density or dynamics of the nucleosomes, coordinated by chromatin remodeling complexes which fall into four categories based on the classification of their ATPase domains: Imitation Switch (ISWI), Chromodomain helicase DNA-binding (CHD), Inositol-Requiring Protein 80 (INO80) and Switching-Defective/Sucrose Non-Fermenting (SWI/SNF) complexes [11–13]. Their common property of DNA translocation provides the force needed to dislodge histone-DNA contacts and is tailored to achieve nucleosome repositioning, sliding, ejection or editing depending on the complex [14]. Chromatin remodeling has been shown to facilitate gene expression in a variety of contexts. For example, INO80 is recruited by Oct4 at pluripotency genes to maintain their accessibility in ES cells [15], and by reducing nucleosome occupancy, it facilitates oncogene transcription in melanoma [16]. Similarly, SWI/SNF remodelers establish accessible enhancers in fibroblasts following their recruitment by lineage-specific transcription factors and FOS/JUN [17], and function to shift nucleosomes away from GATA1 sites in hematopoietic stem cells allowing TAL1-dependent transcription [18]. Conversely, in some cases chromatin remodeling can be inhibitory, such as at the MMLV promoter where SWI/SNF recruitment by the glucocorticoid receptor (GR) subsequently inhibits GR binding [19,20]. These diverse roles are highlighted by the fact that mutations affecting remodeling complexes can both promote and suppress tumor progression [11,21].

Given that enhancer accessibility appears to play a key role in Notch-mediated transcription, we set out to investigate the nucleosome dynamics at target enhancers and to distinguish which chromatin remodelers are critical for enabling target gene activation by Notch. Firstly, we find that Notch signaling regulates nucleosome turnover at target enhancers and promotes incorporation of the histone variant H3.3. Secondly, by testing several classes of chromatin remodelers, we find that the BRM SWI/SNF complex is required for nucleosome turnover and the enhancer accessibility required for the Notch response. Thus, SWI/SNF complexes are vital for the Notch response, and we propose a model whereby dynamic chromatin remodeling poises Notch-responsive genes for rapid activation.

## Results and Discussion

### Ectopic Notch signaling increases the local concentration of histone H3.3 at the E(spl)-C

Previous work has shown that Notch signaling promotes largescale changes in chromatin structure, including rapid increases in histone acetylation and increased chromatin accessibility [4,6–8]. In *Drosophila*, these changes have been most clearly observed at the *Enhancer of split-Complex* (*E(spl)-C*), a 60kb region where 11 highly Notch-responsive genes are concentrated [22–25]. We therefore chose to investigate whether there are any largescale changes in the histone variant H3.3 or canonical histone H3 occupancy at this region following Notch activation, making use of a live tag marking the *E(spl)-C* in *Drosophila* larval salivary glands [8]. Without Notch signaling, both histones H3.3-GFP and H3-GFP were present at low levels at the *E(spl)-C* compared to surrounding regions (Fig 1A and C, Notch-OFF). However, when we activated Notch signaling in this tissue by expressing a constitutively-active form of the Notch receptor, the levels of H3.3-GFP were strongly increased compared to surrounding regions (Fig 1A, Notch-ON). This pattern was found to be reproducible when the relative fluorescence intensity was quantified across the locus in images taken from live salivary glands (Fig 1B). No such change was detected when we examined the effects on histone H3 in a similar manner (Fig 1C and D).

**Figure 1.**
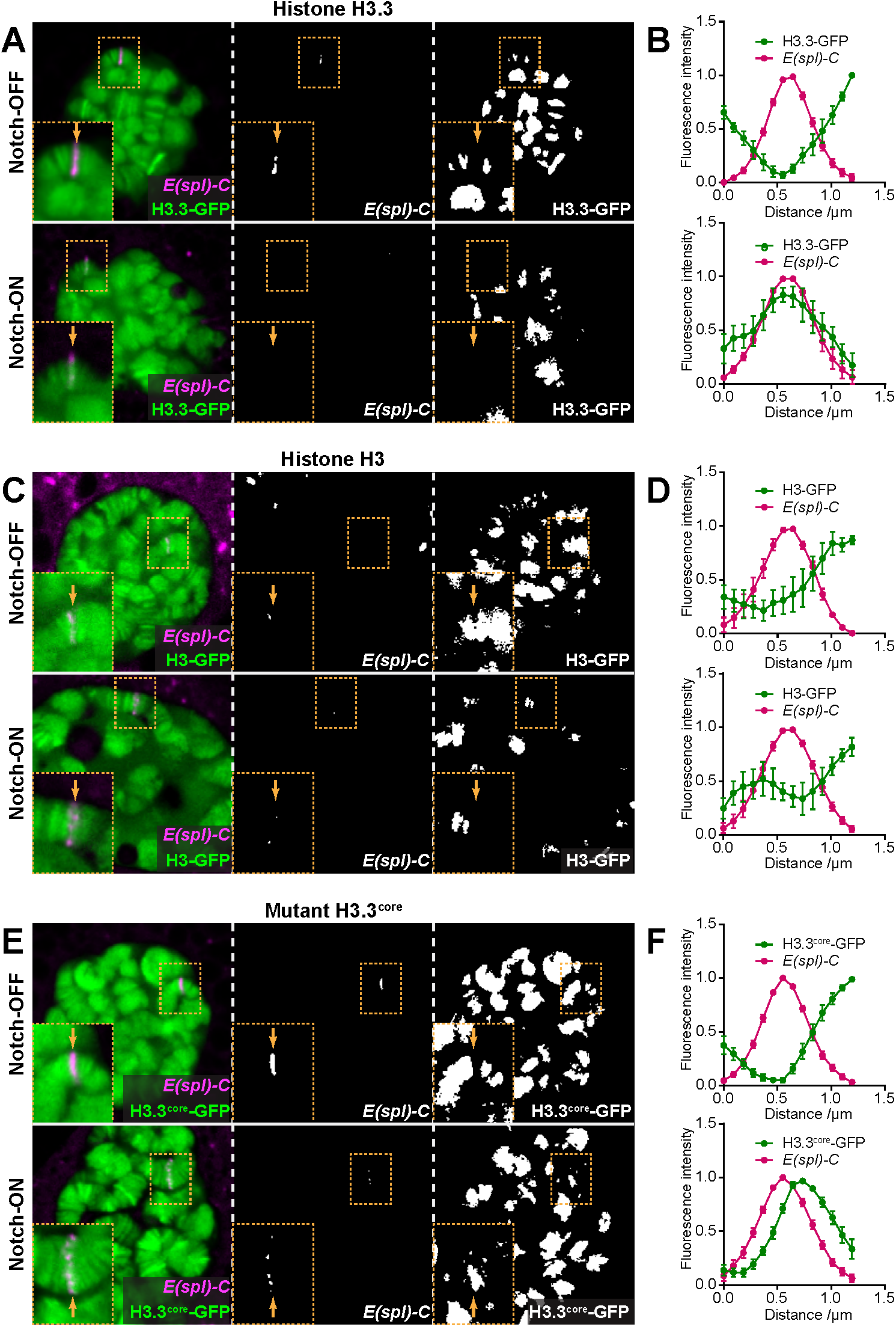
H3.3 levels increase at the *E(spl)-C* in Notch-ON nuclei. A, C, E Live imaging of histone-GFP (green) and ParB-mCherry (magenta) expressed in larval salivary gland nuclei using 1151-Gal4. H3.3-GFP levels are increased at the *E(spl)-C* in the presence of constitutively active Notch, N^ΔECD^ (Notch-ON), compared to control Notch-OFF nuclei expressing LacZ (A). The same is seen with H3.3^core^-GFP (E), but there is little change in H3-GFP between Notch-OFF and Notch-ON nuclei (C). ParB-mCherry binds to its cognate *int* DNA sequence inserted within the *E(spl)-C* [8,48]. Yellow dotted box contains *E(spl)-C* and yellow arrow indicates position of *E(spl)-C* on chromosome. B, D, F Quantifications of relative fluorescence intensity of histone-GFP and ParB-mCherry across the *E(spl)-C* in Notch-OFF (upper) and Notch-ON (lower) conditions. Mean+/-SEM; n >= 5.

Histone H3.3 has been associated with actively-transcribed genes and can be incorporated into the chromatin independently of DNA replication [26,27]. To verify that the Notch-dependent increase in H3.3-GFP was replication-independent, we used a mutant form of histone H3.3, H3.3^core^-GFP, which is only incorporated in a replication-independent manner [26]. Indeed, with H3.3^core^-GFP we saw the same pattern as with H3.3-GFP (Fig 1E and F), suggesting that the local increase in H3.3 concentration at the *E(spl)-C* is not due to an increased level of endoreplication at this locus, and thus likely represents changes associated with Notch-induced transcription.

### The BRM chromatin remodeling complex is required for Notch-responsive accessibility

The incorporation of histone H3.3 at Notch-regulated genes, along with the previously detected changes in accessibility, suggest an involvement of chromatin remodeling complexes and/or histone chaperones. In order to ascertain which of these are needed, we tested if any remodelers or chaperones are required for the recruitment of Su(H) to the *E(spl)-C.* We have previously shown that Notch activity promotes robust recruitment of Su(H)-GFP, detectable as a band of fluorescence when salivary gland nuclei are imaged live (Fig 2A) [8]. We therefore performed RNAi knockdown of different chromatin remodelers and histone chaperones and assessed the impact on Notch-dependent Su(H) recruitment. In the majority of cases we detected little or no change (Fig 2A and B). For example, knockdown of components in the ISWI, NuRD and INO80 complexes failed to perturb Su(H) recruitment (for review of chromatin remodelers in *Drosophila,* see [28]). Likewise, knockdown of chromatin assembly factors or H3.3-specific chaperones such as DEK [29] or Yemanuclein (YEM) [30] had no effect. In contrast, depletion of core components of the BRM (BAF/PBAF) SWI/SNF chromatin remodeling complex [31,32] had a striking effect.

**Figure 2.**
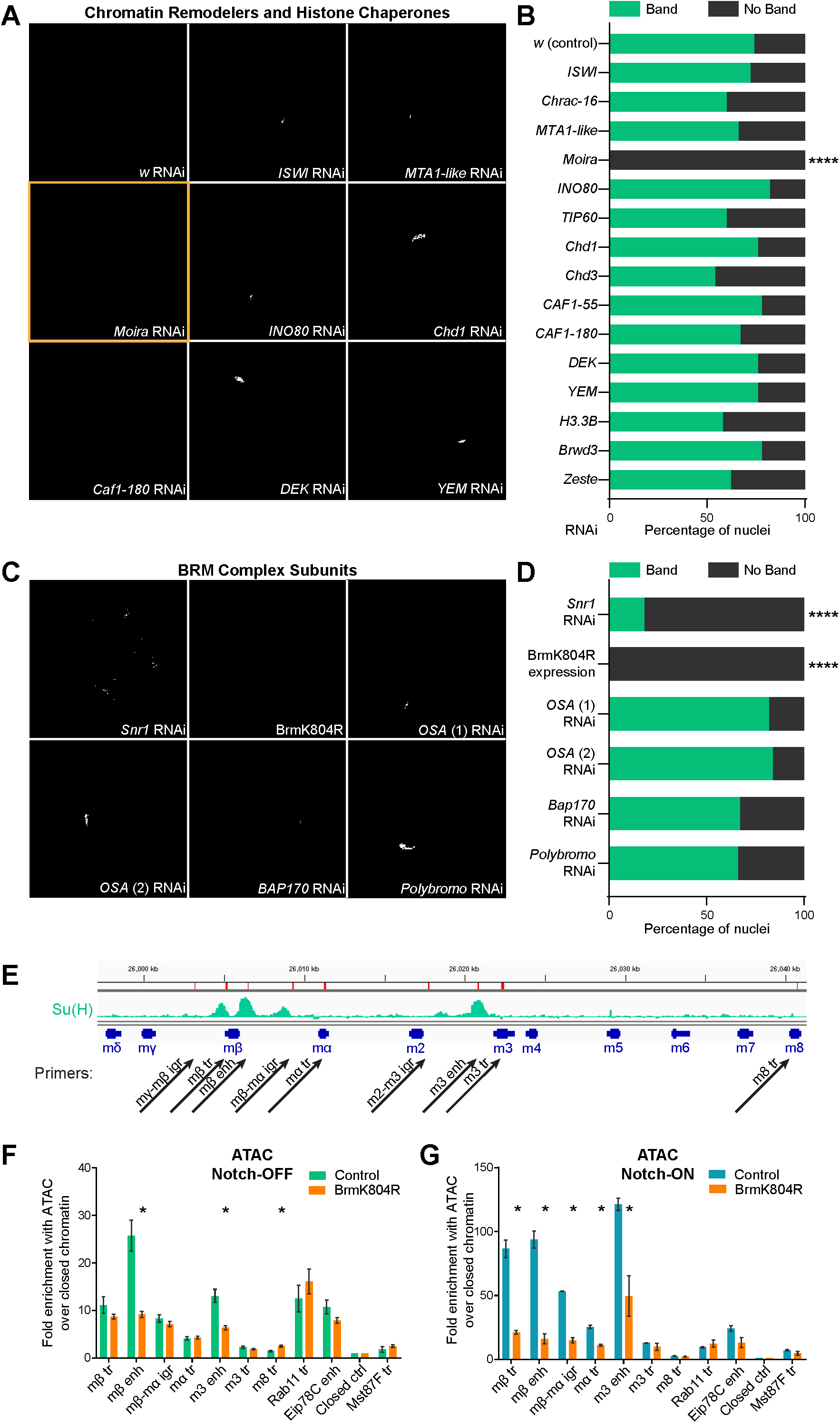
The BRM complex is required for Su(H) recruitment and chromatin accessibility. A, C Effects from depleting chromatin remodelers and histone chaperones, as indicated (wide range shown in A, BRM complex components shown in C), on recruitment of Su(H)-GFP in Notch-ON nuclei (expressing N^ΔECD^). *w* RNAi is a control and BrmK804R is expression of dominant-negative Brm. Different *OSA* RNAi stocks used in C are denoted by (1) and (2). In all conditions except *Moira* RNAi (A), *Snr1* RNAi and BrmK804R expression (C), nuclei exhibit a bright accumulation of Su(H)-GFP at a single locus when imaged live. B, D Percentage of Notch-ON nuclei retaining a single clear band of Su(H)-GFP when the indicated RNAi is co-expressed with N^ΔECD^. **** A significant fraction of nuclei lost the fluorescent band when core components of the BRM complex were perturbed; p<0.0001, two-tailed Fisher’s exact test. E Genomic region encompassing the *E(spl)-C*; green graphs indicate ChIP enrichment for Su(H) in Kc167 cells (Log_2_ scale is-0.5 to 2.9); gene models are depicted in dark blue. Positions of primer pairs used in qPCR experiments are indicated with black arrows. Abbreviations are as follows: “igr” = intergenic region, “tr” = transcribed region and “enh” = enhancer. F, G Chromatin accessibility in Notch-OFF (F) and Notch-ON (G) salivary gland nuclei measured by ATAC-qPCR; fold enrichment at the indicated regions compared to a “closed ctrl” region. Expression of dominant-negative Brm, BrmK804R, led to reduced accessibility of *E(spl)mβ-HLH* and *E(spl)m3-HLH* enhancer regions in Notch-OFF conditions, and to a more widespread reduction in accessibility in Notch-ON conditions. “Eip78C enh” corresponds to the ecdysone receptor-binding region of the Eip78C enhancer, which is highly accessible but not Notch-responsive; “Rab11 tr” and “Mst87F tr” represent highly and lowly-expressed control genes respectively. Mean +/-SEM; n = 3; * p<0.05 with two-tailed Welch’s t-test comparing LacZ and BrmK804R samples.

Knockdown of Moira (SMARCC1/2) eliminated visible recruitment of Su(H)-GFP in all nuclei, while knockdown of Snr1 (SMARCB1) prevented the formation of a single clear band of recruitment in most nuclei (Fig 2A-D). Additionally, expression of a commonly-used dominant negative form of the Brahma ATPase BrmK804R [33] had the same effect as Moira, preventing formation of the Su(H)-GFP band (Fig 2C and D), showing specifically that the ATPase activity of the BRM complex was required.

Two different BRM complexes have been reported, BAP and PBAP (BAF and PBAF), which are distinguished by specific subunits OSA (ARIID1A/B) in BAP or BAP170 (ARID2) and Polybromo (PBRM1) in PBAP [31,32]. Surprisingly, these subunits do not appear to be essential for Notch-dependent Su(H) recruitment. A robust band of Su(H)-GFP is still detectable in nuclei depleted for OSA, BAP170 or Polybromo (Fig 2C and D), even though little or no detectable protein or RNA remains (Fig EV1). This suggests that either the two complexes can compensate for each other or the specialized subunits are not necessary for the Notch-mediated effects on chromatin.

Su(H) recruitment in the Notch-ON condition correlates with increased chromatin accessibility [8]. We therefore used the assay for transposase accessibility (ATAC) [34] to determine whether the BRM complex is required for this Notch-induced change, using qPCR to analyze different regions of the *E(spl)-*C. Expression of BrmK804R had a very localized effect on accessibility measured with ATAC, causing a strong reduction at the *E(spl)mβ-HLH* and *E(spl)m3-HLH* enhancer regions in both Notch-OFF (Fig 2F) and Notch-ON (Fig 2G) conditions. The effects in the Notch-ON condition were the most dramatic, with BrmK804R largely abolishing the increases in accessibility induced by Notch across the *E(spl)-C* so that the locus resembled that in the Notch-OFF condition.

To rule out the possibility that the effects of BrmK804R on accessibility were indirect, resulting from a reduced Su(H) recruitment, we knocked down Su(H) with RNAi in the Notch-OFF condition and performed ATAC. Su(H) RNAi resulted in an increased accessibility across the *E(spl)-*C (Fig EV2), consistent with the known role of Su(H) as a repressor of target genes in the absence of Notch signaling.

Together these results demonstrate that the BRM complex is necessary to maintain a degree of accessibility at enhancers, even before the cells experience Notch signaling, and is then essential for the Notch activity-dependent increase in accessibility of the *E(spl)-C*.

### The BRM complex is required for acute Notch responses in Kc167 cells

To test the role of the BRM complex in a system where we can acutely manipulate Notch activity, we turned to *Drosophila* Kc167 cells. In these cells, Notch signaling is rapidly activated by the addition of the calcium chelator EGTA, which by destabilizing the negative regulatory region, elicits the rapid cleavage of the Notch receptor and activates target genes within 30 minutes [4,24,35]. As in the salivary gland, gene activation is accompanied by an increase in Su(H) recruitment, detectable by chromatin immunoprecipitation (ChIP) [4]. To test the involvement of the BRM complex in this context, we performed RNAi for two core components of the BRM complex, Brm and Snr1, and analyzed the effects on Su(H) recruitment by ChIP with qPCR. Both in Notch-OFF (Fig 3B) and Notch-ON (Fig 3C) cells, the level of Su(H) recruitment was decreased when Brm or Snr1 were depleted by RNAi, showing that the BRM complex is essential for Su(H) recruitment. The transcription of the target genes *E(spl)mβ-HLH* and *E(spl)m3-HLH,* which are usually strongly induced following Notch activation, was also decreased by Brm RNAi (Fig 3D).

**Figure 3.**
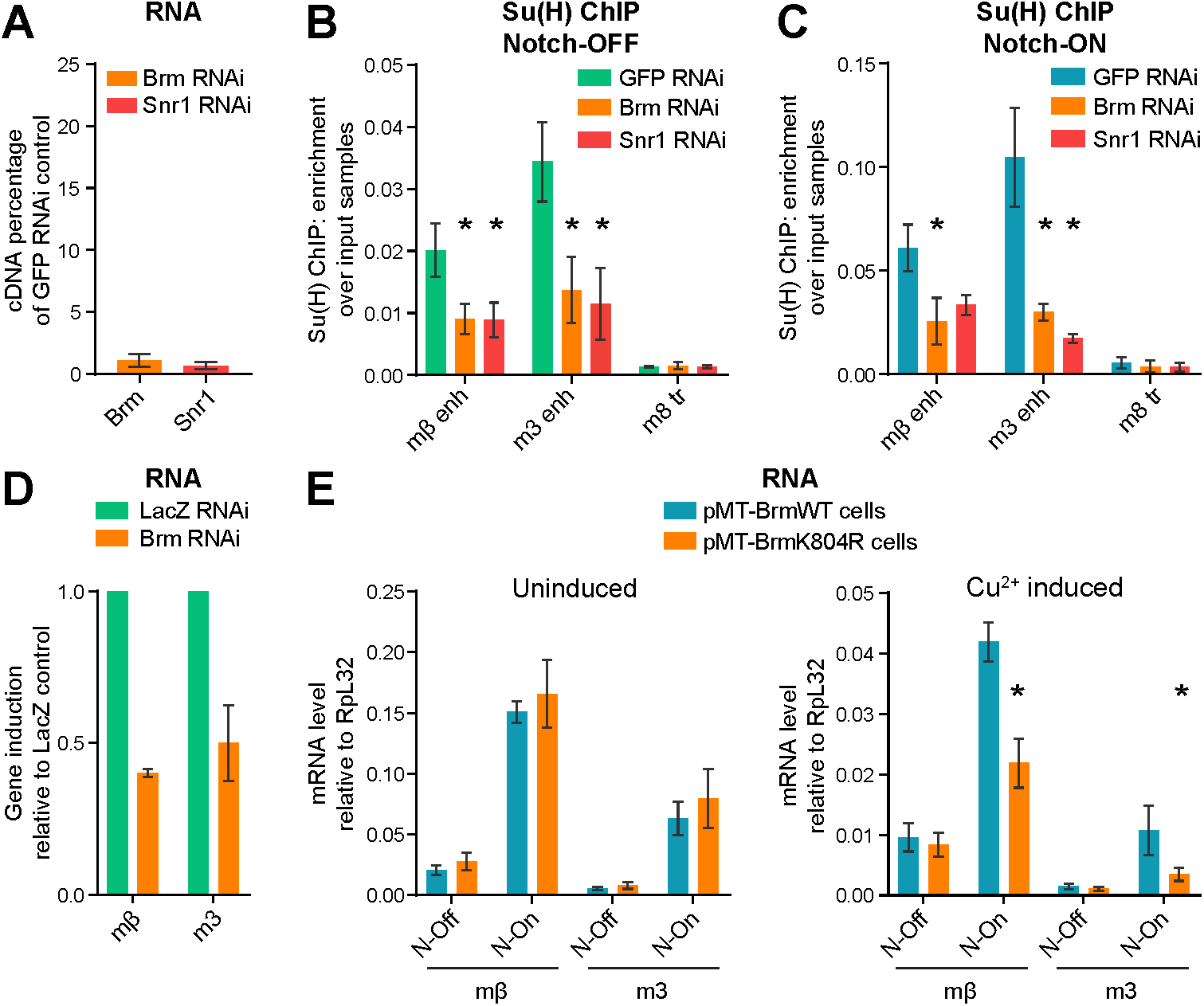
The BRM complex is required for Su(H) recruitment and Notch-dependent transcription in Kc167 cells. A Effect of *Brm* and *Snr1* RNAi on *Brm* and *Snr1* cDNA levels respectively, measured by reverse transcription-qPCR; percentage cDNA compared to *GFP* RNAi. The knockdowns are highly effective, with only 1-2% of *Brm* and *Snr1* cDNA remaining detectable. B, C Knockdown of components of the BRM complex reduces Su(H) recruitment both in Notch-OFF (B) and Notch-ON (C) conditions. Fold enrichment of Su(H) occupancy at the indicated positions detected by ChIP, relative to input, in Kc167 cells treated with *Brm, Snr1* or *GFP* RNAi as a control. Notch-ON conditions (C) were induced by 30 minutes of EGTA treatment. Mean+/-SEM; n = 3 (B) and 2 (C); * p<0.05 with one-tailed student’s t-test compared to *GFP* RNAi control. D Effect of *Brm* RNAi on *E(spl)mβ-HLH* and *E(spl)m3-HLH* induction by Notch activation (EGTA treatment) measured by reverse transcription-qPCR; shown as fold difference to *LacZ* RNAi control. Mean +/-SEM; n = 3. E Effects of Brm dominant-negative on expression of *E(spl)mβ-HLH* and *E(spl)m3-HLH* measured by reverse transcription-qPCR. Expression was analyzed in stable cell lines containing pMT-inducible BrmWT or BrmK804R in the absence (left, uninduced) or presence of copper sulfate (right, Cu^2+^ induced). The response of *E(spl)mβ-HLH* and *E(spl)m3-HLH* to Notch activation (“N-On” = EGTA treatment vs. “N-Off” = PBS control) was reduced in the BrmK804R-expressing cells compared to BrmWT-expressing cells, only when induced with copper (right graph). Mean +/-SEM; n = 2 (left) and 3 (right); * p<0.05 with one-tailed student’s t-test comparing BrmWT and BrmK804R.

In order to confirm that the ATPase activity of the BRM complex was essential, we made stable cell lines expressing BrmK804R or the wild type form, BrmWT as a control (under control of the copper-inducible pMT promoter) [36]. Expression of *E(spl)mβ-HLH* and *E(spl)m3-HLH* was rapidly induced by Notch activation in control conditions (Fig 3E, left). However, following copper-induced expression of BrmK804R, cells had a significantly reduced upregulation of *E(spl)mβ-HLH* and *E(spl)m3-HLH* compared to cells expressing BrmWT (Fig 3E, right). This shows that the ATPase function of the BRM complex is key to the Notch response in these cells.

### Nucleosome turnover increases with Notch signaling and is dependent on the BRM complex

Chromatin remodelers are thought to slide, replace or eject nucleosomes [13]. Even the short pulse of activity in Kc167 cells was sufficient to bring about a change in chromatin accessibility measured with ATAC (Fig EV3), suggesting that the BRM complex could be moving or depleting histones at the Notch-regulated enhancers. Additionally, the histone variant H3.3 has been associated with nucleosome turnover [37,38]. Given the results showing changes in accessibility and histone H3.3 levels, we were prompted to measure whether nucleosome turnover is occurring. To do this we used the CATCH-IT technique, which relies on the incorporation of a methionine analog called azidohomoalanine into newly-synthesized proteins [39,40]. Click chemistry is used to biotinylate this residue so that any chromatin containing newly-synthesized histones can therefore be isolated. We performed CATCH-IT in Kc167 cells, incubating them with media containing azidohomoalanine after a period of methionine starvation, in the presence or absence of NICD. To achieve this, we used a cell line where NICD is expressed from the copper-inducible pMT promoter [4]. Using this approach we detected differential levels of histone turnover. Notably, the Su(H)-binding enhancer regions had high levels of nucleosome turnover compared to surrounding regions, and this turnover was significantly increased in the presence of NICD (Fig 4A).

**Figure 4.**
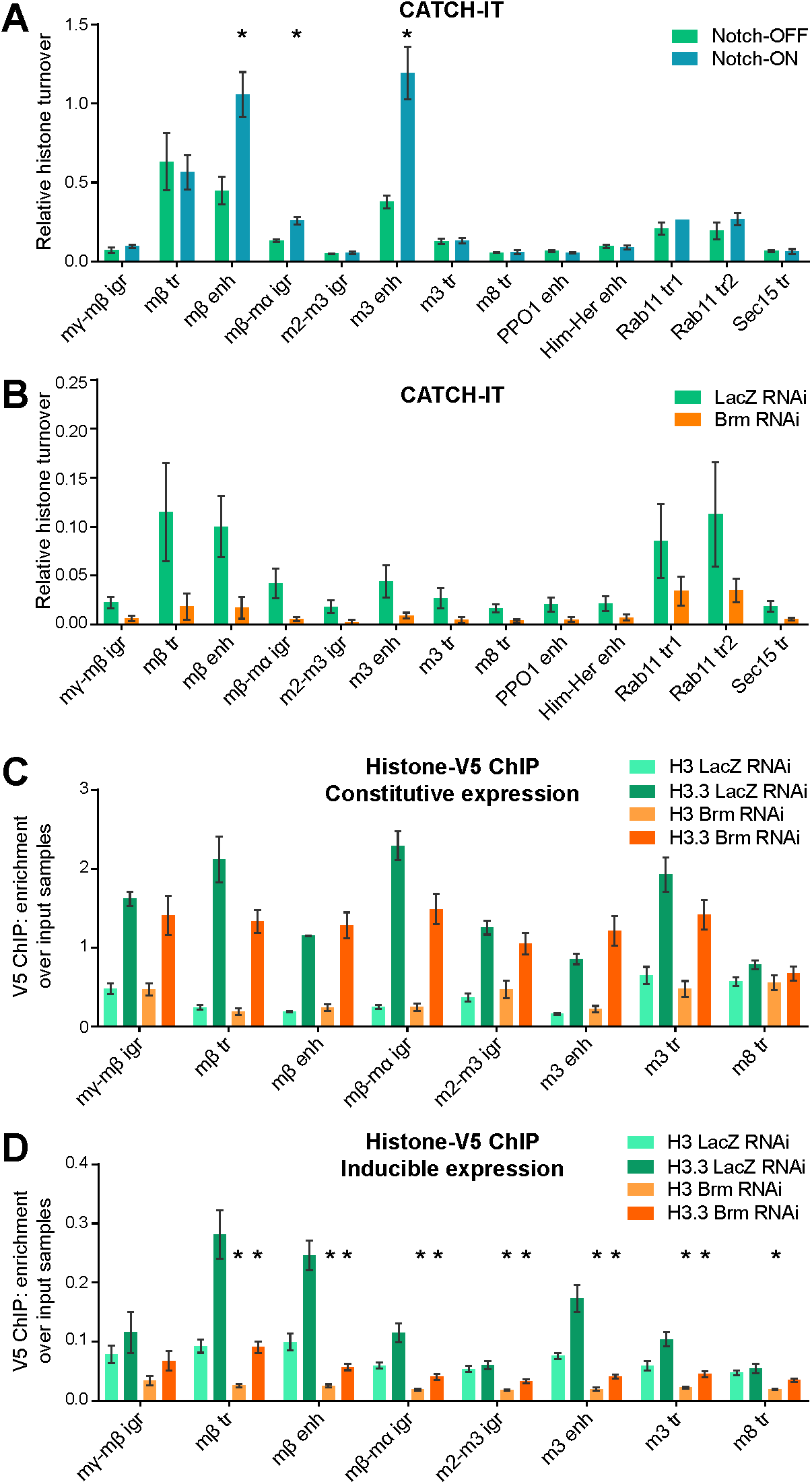
Nucleosome turnover at Su(H)-bound enhancers is increased in Notch-ON cells and is dependent on the BRM complex. A Nucleosome turnover measured by CATCH-IT-qPCR; fold enrichment over input samples after methionine control level subtraction. Su(H)-bound enhancers show increased nucleosome turnover in response to Notch signaling. Notch signaling is activated in Kc167 cells by 6 hours of copper induction of pMT-NICD with copper excluded in the control. Positions of *E(spl)-C* primers are shown in Fig 2E; the remaining primers are control non-Notch-responsive regions. Mean +/-SEM; n = 3 (primers m?-mβ igr to Sav tr) or 2 (primers PPO1 enh to Sec15 tr); * p<0.05 with two-tailed Welch’s t-test comparing with and without copper induction. B *Brm* RNAi reduces nucleosome turnover. CATCH-IT-qPCR results as in A after *Brm* or*LacZ* RNAi as a control. Mean +/-SEM; n = 3. C Overall enrichment of H3.3 over H3 across the *E(spl)-C*. V5 ChIP-qPCR in Kc cells expressing H3-V5 or H3.3-V5 from a ubiquitous promoter after *LacZ* or *Brm* RNAi treatment, shown as fold enrichment over input samples. H3-V5 and H3.3-V5 show a differential pattern across the *E(spl)-C* but changes caused by *Brm* RNAi are minimal. Mean +/-SEM; n = 3. D *Brm* RNAi reduces incorporation of H3.3. V5 ChIP-qPCR in Kc cells after *LacZ* or *Brm* RNAi treatment in cells with H3-V5 or H3.3-V5 expression induced from the pMT promoter by 3 hours of copper treatment, shown as fold enrichment over input samples. Mean +/-SEM; n = 3. * p<0.05 with two-tailed Welch’s t-test compared to *LacZ* RNAi control.

We then tested whether knockdown of the BRM complex would affect the levels of nucleosome turnover measured with CATCH-IT. Depletion of Brm by RNAi resulted in a consistent decrease in histone turnover, demonstrating that the BRM complex has a critical role in the process of nucleosome turnover (Fig 4B).

Given that we observed increased histone H3.3 recruitment in Notch-ON cells *in vivo,* we next sought to measure the effects on the histone variant H3.3 by expressing V5-tagged histone proteins and performing ChIP [41]. When H3-V5 and H3.3-V5 were expressed from a constitutive promoter, it was evident that H3.3 predominates over H3 throughout the Notch-responsive regions of the *E(spl)-*C, while the distal region at the *E(spl)m8-HLH* gene had similar levels of both variants (Fig 4C). Neither Notch activity (Fig EV4) nor depletion of Brm (Fig 4C) had a large impact on these distributions of H3 or H3.3, arguing that there is no gross change in the overall levels of histones H3 or H3.3 during an acute Notch response, and that the BRM complex is not essential for the incorporation of H3.3 per se.

To test the dynamics of the two histone variants in this system, we then expressed H3-V5 and H3.3-V5 from the pMT promoter so that we could monitor their incorporation over a short timescale. By approximately 90 minutes after their induction, the labelled histones had started to be incorporated into the chromatin (Fig EV5). By three hours after induction, differential incorporation of H3.3-V5 could be observed at specific regions, replicating the pattern seen across the *E(spl)-C* with CATCH-IT, while the levels of H3-V5 incorporation were lower and largely uniform. Crucially, the incorporation of H3.3-V5 was greatly reduced following depletion of Brm, in agreement with the BRM complex having a key role in nucleosome turnover (Fig 4D).

In summary, we have shown that the BRM chromatin remodeling complex is essential for Notch-responsive gene activation. We propose that the BRM complex is required to maintain high levels of accessibility at Notch-responsive enhancers where it promotes rapid nucleosome turnover. This has two implications for Notch signaling. Firstly, the BRM complex is important for the local turnover of nucleosomes that enables Su(H) to access enhancers with its co-repressors in the absence of Notch signaling, poising the enhancers for activation (Fig 5A). Secondly, it is responsible for the dramatic increase in chromatin accessibility at responsive genes following Notch activation (Fig 5B). It is possible that the BRM complex also plays a key role in switching off the Notch response upon cessation of signaling, since the continual turnover of nucleosomes provides a mechanism for the rapid switching of chromatin states.

**Figure 5.**
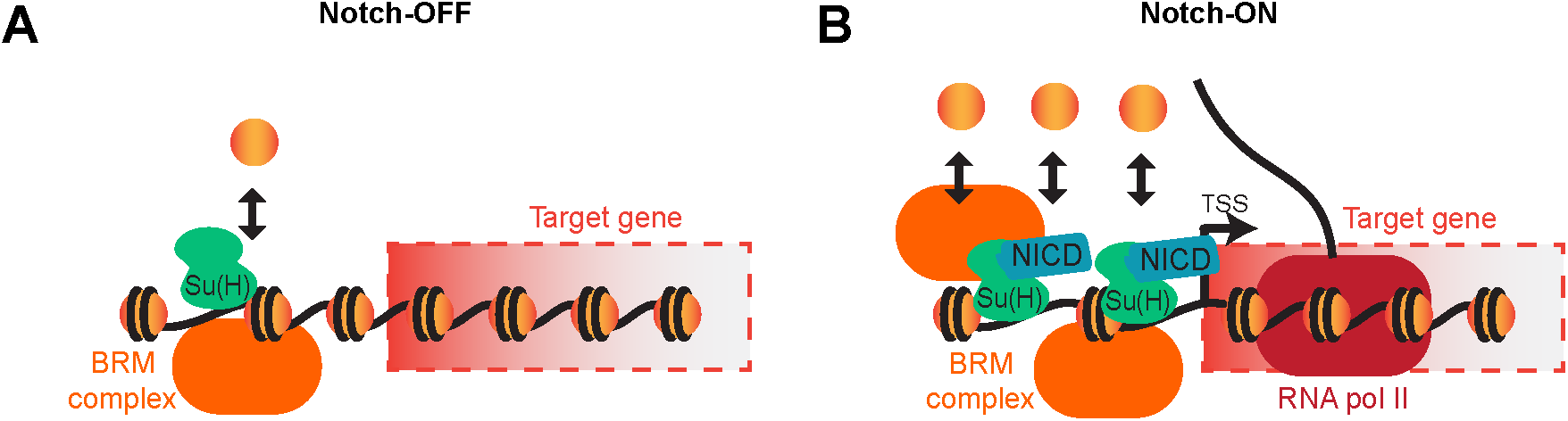
Model of BRM complex action. A In the absence of Notch signaling, the BRM complex maintains the accessibility of Notch-responsive enhancers to allow Su(H) recruitment by promoting nucleosome turnover. B When Notch signaling is activated, the nucleosome turnover at Notch-responsive enhancers increases, increasing the accessibility of the chromatin and allowing more Su(H) to bind. The BRM complex is essential for the process to occur. Target genes are activated via co-activators.

The use of techniques measuring the dynamic nature of nucleosomes have allowed us to uncover the BRM complex-dependent turnover, since there is little change in the steady-state levels of histones bound to the DNA. The histone variant H3.3 more readily undergoes DNA replication-independent turnover than the canonical histone H3 and therefore its incorporation is more sensitive to BRM complex depletion. However, the non-essential function of histone H3.3 [42] would suggest that histone H3 can compensate in its absence and that the BRM complex does not act exclusively on histone H3.3 as a substrate.

Our data provide new evidence for the involvement of SWI/SNF chromatin remodeling in Notch signaling and give mechanistic insight into the relationship between the two. The results are fully consistent with a previous observation that BRM was recruited to Hes1 and Hes5 in mouse myoblasts before Notch induction [43], and with genetic data that hinted at a role for the BRM complex in regulating Notch target genes [44]. Thus, it is likely that the recruitment of SWI/SNF complexes will be a key step in selecting enhancer repertoires in different contexts. A similar model has been proposed for serum stimulation acting via FOS/JUN, where serum-induced changes in chromatin accessibility relied on BRM recruitment [17]. The SWI/SNF-dependent nucleosome turnover is therefore likely to have an integral role in generating accessible enhancer landscapes critical for the specificity of signaling pathway responses.

## Materials and Methods

### Fly stocks

For expression of all UAS constructs in the salivary gland, 1151-Gal4 was used (L S Shashidhara, Centre for Cellular and Molecular Biology, Hyderabad, India) [45]. Notch signaling was activated by UAS-N^ΔECD^ [46,47]. The *E(spl)-C* was imaged using the ParB-INT DNA tagging system where UAS-ParB1-mCherry was expressed in the presence of the INT sequence inserted between *E(spl)m7-HLH* and *E(spl)m8-HLH* [8,48]. Histone-GFP imaging made use of UAS-H3-GFP, UAS-H3.3-GFP and UAS-H3.3^core^-GFP constructs (flies kindly provided by Kami Ahmad) [26,49]. Su(H) recruitment was monitored using Su(H)-GFP [8]. Dominant negative Brm was expressed from UAS-BrmK804R [33]. RNAi lines used are listed in Table 1.

**Table 1.**
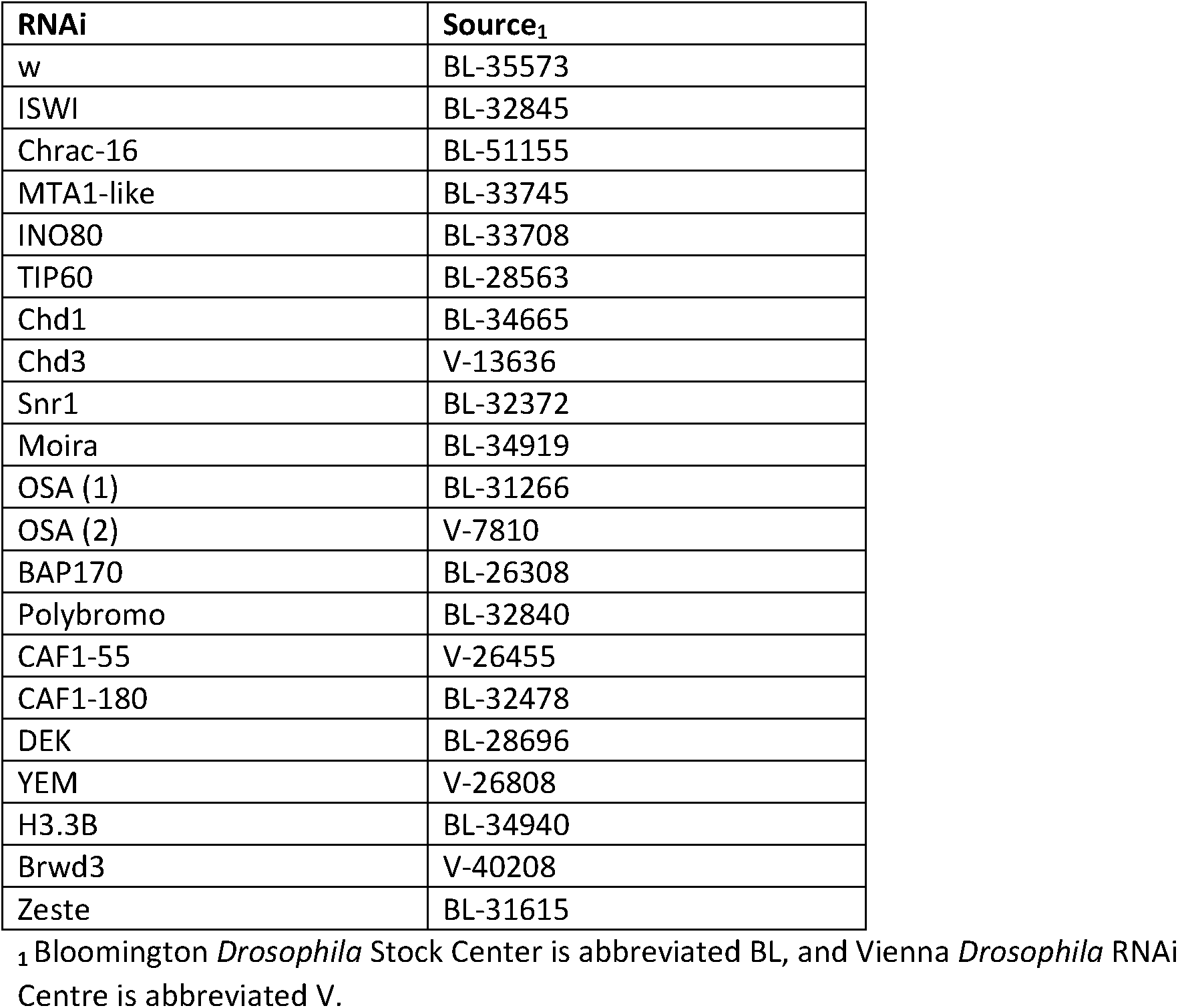
RNAi lines used.

### Live imaging of salivary gland nuclei

Salivary glands were dissected and mounted as described previously [8], using Shields and Sang M3 Insect Medium (Sigma S3652) supplemented with 5% FBS (Sigma F9665) and 1x Antibiotic-Antimycotic (Gibco 15240062) for dissection and the same medium with the addition of 2.5% methyl-cellulose (Sigma) for mounting. For DNA stains, salivary glands were incubated in dissecting media containing 200 µg/mL Hoechst 33342 (ThermoFisher) for 10 minutes at room temperature before washing with PBS and mounting.

Image acquisition was performed with Nikon D-Eclipse C1 confocal microscope using lasers at 405, 488 and 543 nm. Images captured of nuclei used the 60x oil objective with a 4.5x zoom level and images of glands used the 40x oil objective. To monitor Su(H)-GFP recruitment, nuclei were scanned slowly through the Z-stack using a 2x zoom level while looking for accumulations of fluorescence. 10 glands and five nuclei per gland were analyzed and scored per condition, with the five nuclei closest to the coverslip chosen each time.

### Immunofluorescence staining

Staining of salivary glands was performed as described [8] except for the following changes. Glands were permeabilized with 1% Triton X-100 in PBS for 30 minutes. Antibodies against OSA and BAP170 were gifts from Peter Verrijzer [50] and were used at dilutions of 1:200 and 1:100 respectively.

### Kc167 cell culture, Notch activation and generation of stable lines

Kc167 cells (Drosophila Genomics Resource Center) were cultured in Schneider’s *Drosophila* medium (Gibco 21720024) supplemented with 5% FBS (Sigma F9665) and 1x Antibiotic-Antimycotic (Gibco 15240062) at 25°C. Notch was activated either by NICD expression from the pMT vector (cell line described below) or by EGTA treatment where media was replaced with 4 mM EGTA (Bioworld) in PBS for 30 minutes.

Stable cell lines were generated by transfection followed by antibiotic selection. 18 µg of the relevant plasmid was mixed with 925 µL Opti-MEM (Gibco 31985070) and 54 µL FuGENE HD Transfection Reagent (Promega E2311) at room temperature for 30 minutes before adding dropwise to cells plated in 10 cm plates. After 24 to 48 hours media was replaced to contain antibiotic selection. Cells were grown in the presence of antibiotic and experiments were performed after significant cell death and recovery had taken place to indicate selection (usually after approximately 3 weeks).

CATCH-IT was performed in the pMT-NICD cell line generated previously [4], where cells were maintained with 2 µg/mL puromycin (Sigma).

Cell lines expressing BrmWT and BrmK804R were generated using plasmids kindly provided by Neus Visa [36]. BrmK804R was re-made by mutagenesis to ensure homogeneity between the two constructs using Pfu polymerase with the primers listed in Table 2. The BrmWT and BrmK804R sequences were then cloned into the pMT-puro vector (Addgene 17923) by digestion with SpeI and PmeI (NEB) and ligation (T4 ligase; Promega). After transfection of pMT-BrmWT and pMT-BrmK804R, cells were selected with 5 µg/mL and maintained with 2 µg/mL puromycin.

**Table 2.**
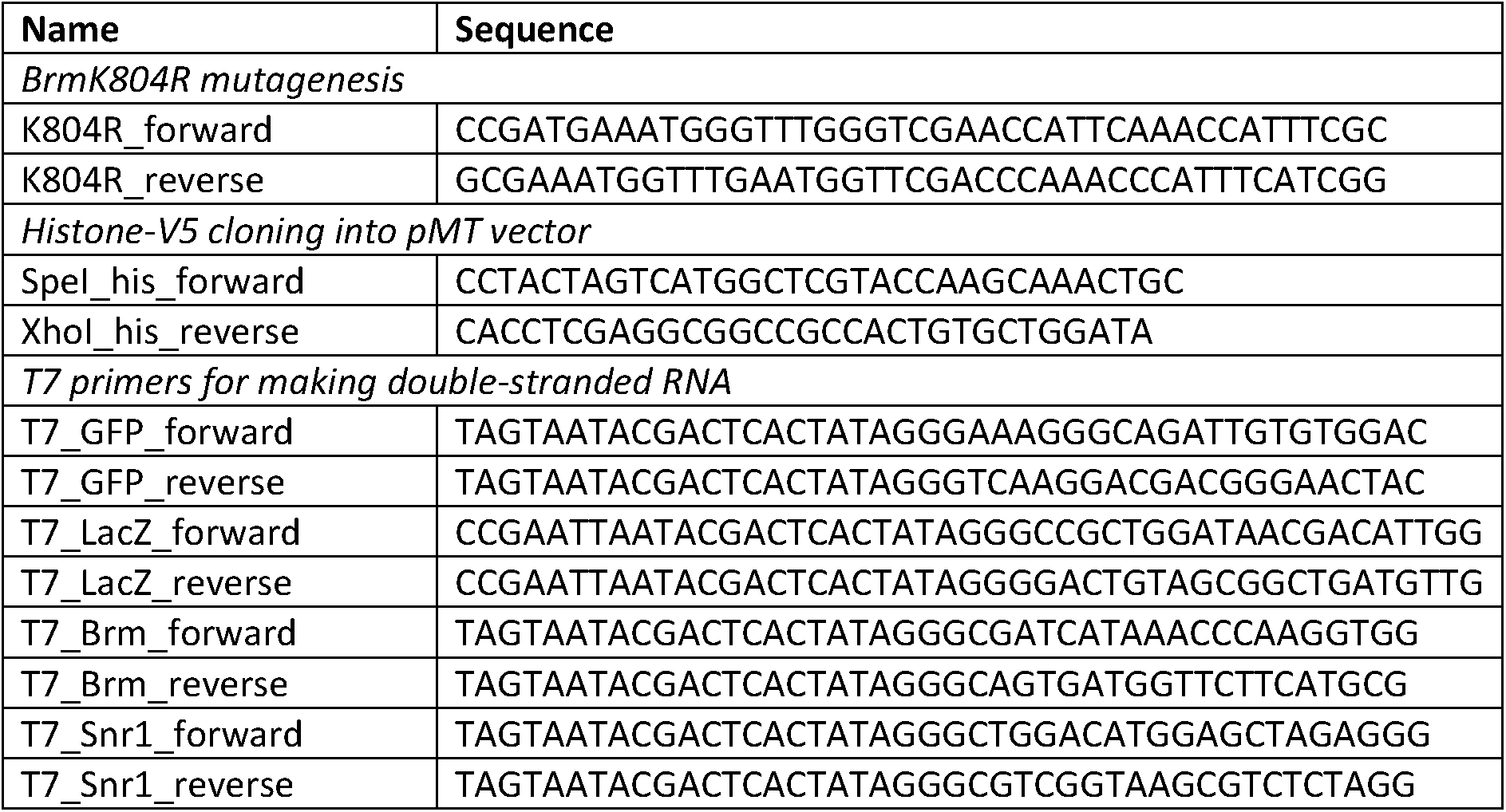
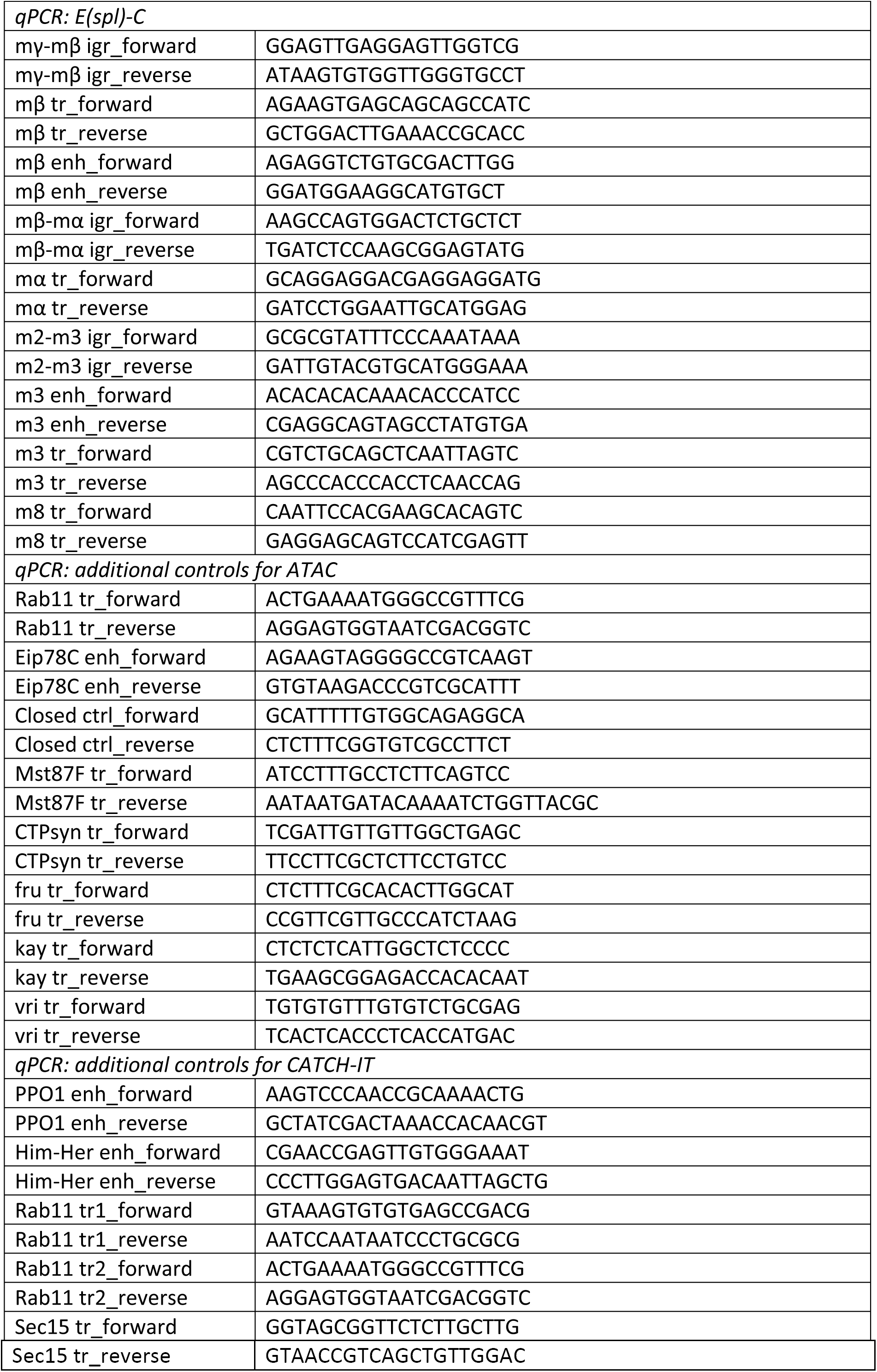
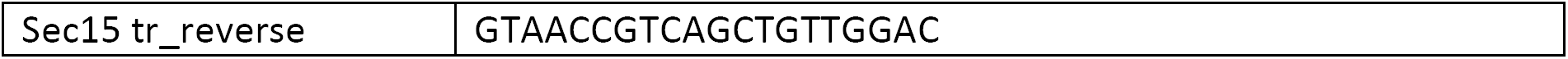
All primers used.

Constitutive expression of histone-V5 proteins made use of pIB-H3-V5 and pIB-H3.3-V5 plasmids kindly provided by Dirk SchÜbeler and used as described [41]. Cells were selected with 50 µg/mL and maintained with 20 µg/mL blasticidin (ThermoFisher R21001). For Notch activation in these cells, they were further transfected with pMT-NICD and selected then maintained with 5 µg/mL and 2 µg/mL puromycin respectively.

For inducible expression of histone-V5 proteins, H3 and H3.3 sequences were cloned from pIB-H3-V5 and pIB-H3.3-V5 into the pMT-puro vector using SpeI and XhoI sites (NEB) with the primers listed in Table 2. After transfection of pMT-H3-V5 and pMT-H3.3-V5, cells were selected with 5 µg/mL and maintained with 2 µg/mL puromycin.

To induce expression from all pMT constructs, 5mM CuSO_4_ was added to normal culture media. Induction was performed for 24 hours for experiments with pMT-BrmWT and pMT-BrmK804R, and for the lengths of time specified for other experiments (see CATCH-IT method for details in this experiment).

### Assay for transposase-accessible chromatin

ATAC using salivary glands was performed exactly as described previously with no changes [8]. ATAC was performed in Kc167 cells in a similar manner with the following changes. After a 30-minute EGTA treatment in 10 cm culture plates containing approximately 40 million cells, cells were immediately harvested taking a quarter of the cells for the experiment (roughly 10 million). Cells were pelleted at 500xg, 4°C for 5 minutes, washed in 10 mL of cold PBS and pelleted again. The cells were then lysed by resuspending in 50 µL lysis buffer (10 mM Tris-HCl, pH7.4, 10 mM NaCl, 3 mM MgCl_2_, 0.3% NP-40), vortexing for 10 seconds, keeping on ice for 3 minutes, and vortexing again. Nuclei were pelleted at 400xg, 4°C for 5 minutes and resuspended in 30 µL TD buffer (Illumina FC-121-1030). 25 µL was used for the tagmentation reaction and the rest of the protocol performed exactly as described previously for salivary glands [8].

### RNAi in Kc167 cells

300 to 800 base-pair regions of *Brm* and *Snr1* DNA were amplified from genomic DNA, with *GFP* or *LacZ* sequences amplified from plasmids as controls, using either Q5 or Phusion High-Fidelity DNA Polymerases (NEB M0491 and M0530 respectively) and overhanging primers containing the T7 promoter sequence listed in Table 2. *In vitro* transcription was performed using the MEGAscript T7 Transcription Kit (Invitrogen AM1334). RNA was purified by phenol-chloroform extraction and then annealed to form double-stranded RNA by heating to 75°C and cooling slowly. 100 µg double-stranded RNA was mixed with 3.5 mL Opti-MEM (Gibco 31985070) and added to approximately 10 million cells in a 10 cm plate for 30 minutes before topping up to 10 mL with normal culture medium. Volumes were scaled down for some smaller experiments.

### RNA extraction and reverse transcription

To extract RNA from Kc167 cells, TRI reagent solution (Invitrogen AM6738) was used followed by phenol chloroform extraction and isopropanol precipitation at-20°C overnight. For reverse transcription, RNA was resuspended in water and first DNase-treated with the DNA-free DNA Removal Kit (Invitrogen AM1906), before reverse transcribing with M-MLV Reverse Transcriptase (Promega M1705) using Oligo(dT)_15_ Primers (Promega C1101). cDNA was diluted 5-fold before analysis with qPCR.

The same protocol was used to extract RNA from salivary glands, exactly as described previously [8].

### Western blot

Approximately 20 million Kc167 cells were lysed in 100 µL lysis buffer (50 mM Tris-HCl, pH8.0, 150 mM NaCl, 10% glycerol, 0.5% triton X-100) on ice for 30 minutes before debris was removed by centrifugation at 13,000xg, 4°C for 30 minutes. Samples were then combined with 2x loading buffer (10 mM Tris-HCl, pH6.8, 20% glycerol, 4% SDS, 0.025% bromophenol blue, 2% mercaptoethanol) and boiled. Proteins were resolved using standard protocols with 15% SDS-PAGE and transferred to nitrocellulose. Blots were probed with antibodies against histone H3 (Abcam ab1791) and V5 (Invitrogen R960-25) at dilutions of 1:1000 and 1:4000 respectively. Horseradish peroxidase-conjugated secondary antibodies were used and detected with the ECL system (GE Life Sciences).

### Chromatin immunoprecipitation

Su(H) and V5 ChIP were performed largely as described previously [4,51], using 2.5 µg goat Su(H) antibody (Santa Cruz Biotechnology, no longer available) and 1-2 µg V5 antibody (Invitrogen R960-25). Briefly, cells were cross-linked with 1% formaldehyde (Sigma F8775) in PBS for 10 minutes at 25°C. After lysis, chromatin was diluted 2-fold for sonication and then a further 5-fold for pre-clearing with goat or mouse IgG and 40 µL protein G or protein A/G PLUS-Agarose (Santa Cruz Biotechnology sc-2002 and sc-2003) for Su(H) and V5 ChIP respectively. Immunoprecipitation was performed with 40 µL of the same beads at 4°C overnight, followed by washes, elution by vortexing, de-crosslinking with 0.3 M NaCl, 0.1 mg/mL RNase A and 0.1 mg/mL proteinase K treatment. DNA was purified with the QIAquick PCR purification kit (Qiagen) and eluted in 100 µL water for analysis with qPCR.

### CATCH-IT

Schneider’s Drosophila medium without methionine (PAN Biotech), supplemented with 5% FBS and 1x Antibiotic-Antimycotic was added to cells for 1 hour, followed by adding either 4 mM azidohomoalanine (Aha; AnaSpec AS-63669) or 4 mM methionine (Sigma) as a control for 4 hours. To activate Notch, pMT-NICD cells were induced with 5 mM CuSO_4_ for 1 hour before the medium was substituted for methionine-free medium, also containing 5 mM CuSO_4_, so that cells were incubated with CuSO_4_ for a total of 6 hours.

CATCH-IT was performed as previously described [40], except where stated otherwise. Briefly, cells were harvested and nuclei were extracted with 30 µL of 10% NP-40. Nuclei were resuspended in 180 µL of HB125 buffer, and the following were added: 5 µL of 2 nM biotin-alkyne (Invitrogen B10185), 10 µL of 100mM THPTA (Sigma 762342) premixed with 2 µL of 100 mM copper sulfate (Jena Bioscience CLK-M1004), and 6 µL of freshly-prepared 500 mM sodium ascorbate (Jena Bioscience CLK-M1005). Cycloaddition reaction was performed for 30 minutes at room temperature on a rotor. Reaction with MNase (Sigma N3755) was performed at 37°C for 3 minutes. After capture with Dynabeads M-280 Streptavidin (Invitrogen 11205) as described, captured chromatin and input chromatin samples were treated with 0.25 mg/mL RNase A (Roche) and 0.25 mg/mL proteinase K (ThermoFisher). DNA was purified with the QIAquick PCR purification kit (Qiagen) and analyzed by qPCR.

### qPCR

All qPCR was performed using LightCycler 480 SYBR Green I Mastermix (Roche 04707516001) as described previously [8]. For reverse transcription experiments, relative amounts of the genes of interest were normalized to the control gene RpL32. For ChIP, immunoprecipitated samples were normalized to input samples. For CATCH-IT, pulldown samples were normalized to input samples and methionine controls were subtracted. All primers used are shown in Table 2.

## Acknowledgements

We thank Jin Li, Damiano Porcelli and members of the Bray Lab for technical advice and helpful discussions. This work was supported by a Programme Grant from the Medical Research Council to SJB (MR/L007177/1) and by a studentship from BBSRC for ZP (1502069).

## Author Contributions

SJB and ZP conceived the project, SJB supervised the work, ZP performed the experiments and analyzed the data, ZP and SJB wrote the manuscript.

## Expanded View Figure Legends

**Figure E1.**
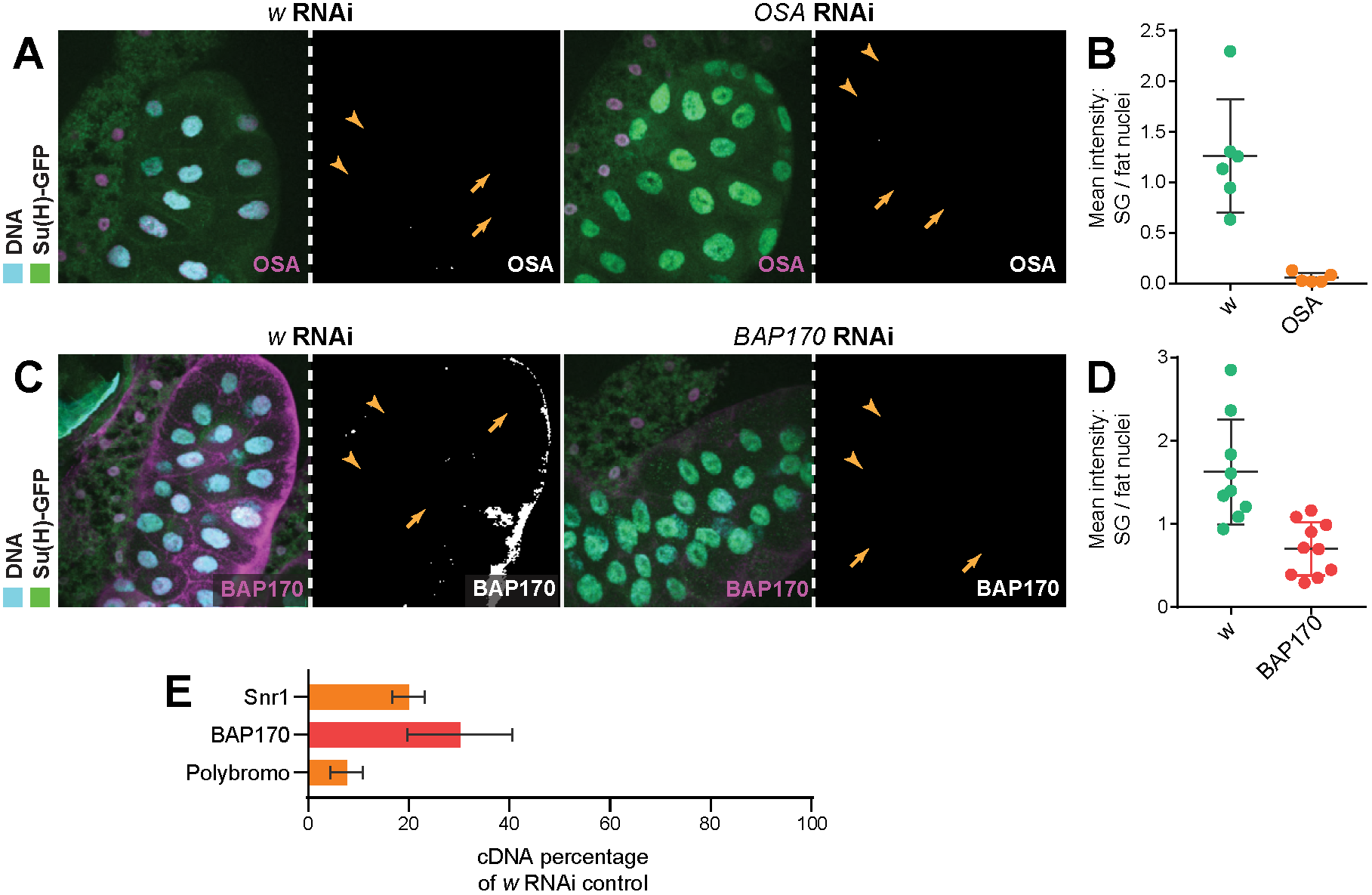
OSA, BAP170 and Polybromo RNAi successfully reduce protein and RNA levels. A, C Immunofluorescence staining of OSA (A; magenta) and BAP170 (B; magenta) in salivary glands expressing *OSA* (stock (2) in Fig 2) and *BAP170* RNAi respectively, compared to *w* RNAi control glands. *OSA* RNAi depletes all detectable OSA protein and *BAP170* RNAi removes most BAP170 protein. Yellow arrows indicate salivary gland nuclei and yellow arrowheads indicate fat cell nuclei for comparison where RNAi is not expressed. B, D Quantifications of OSA (B) and BAP170 (D) nuclear levels from maximum projection images with salivary gland nuclei normalized to fat cell nuclei. E Effect of *Snr1, BAP170* and *Polybromo* RNAi expression in salivary glands on *Snr1, BAP170* and *Polybromo* cDNA levels respectively, measured by reverse transcription-qPCR; percentage cDNA compared to *w* RNAi control. All reduce their respective cDNA levels, with *Polybromo* RNAi causing a greater reduction than *Snr1*, despite not having an effect on Su(H) recruitment.

**Figure E2.**
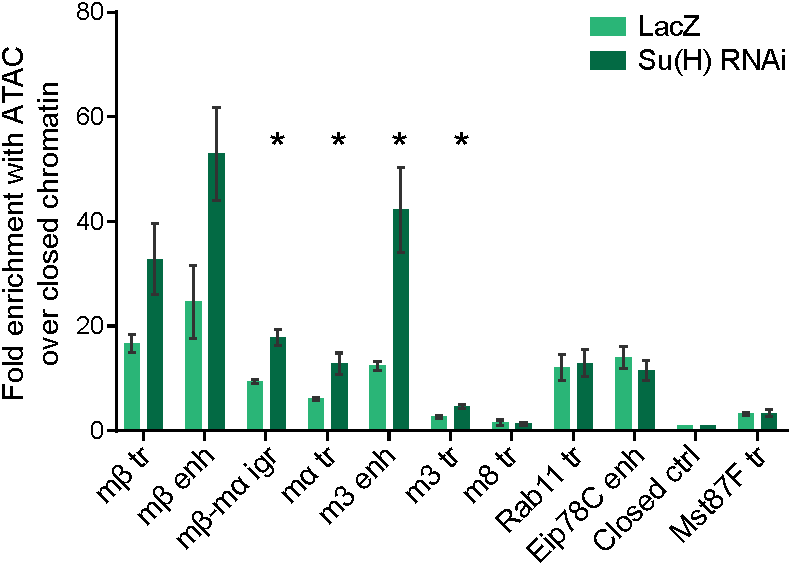
Su(H) depletion increases accessibility across the *E(spl)-C.* Chromatin accessibility in salivary gland nuclei depleted for Su(H) by RNAi, measured by ATAC-qPCR; fold enrichment at the indicated regions compared to a “closed ctrl” region. Accessibility is increased across most of the *E(spl)-C* compared to controls expressing LacZ. Control primer regions are the same as in Fig 2F and G. Mean +/-SEM; n = 3; * p<0.05 with two-tailed Welch’s t-test compared to LacZ controls.

**Figure E3.**
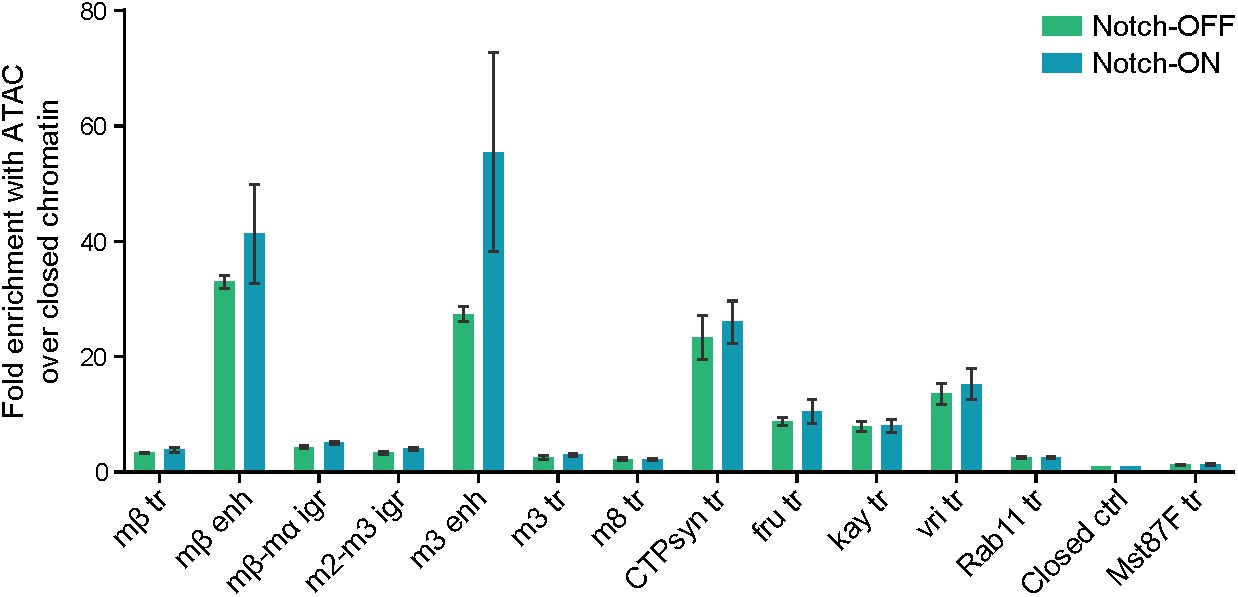
An acute Notch response in Kc167 cells involves increased enhancer accessibility. Chromatin accessibility across the *E(spl)-C* in Notch-ON (EGTA-treated) and Notch-OFF (PBS control) cells detected by ATAC-qPCR. Fold enrichment of the indicated regions compared to a “closed ctrl” region; positions of *E(spl)-C* primers in the genome are shown in Fig 2E. “CTPsyn tr”, “fru tr”, “kay tr” and “vri tr” are highly accessible control regions which do not respond to Notch. Mean +/-SEM; n = 3.

**Figure E4.**
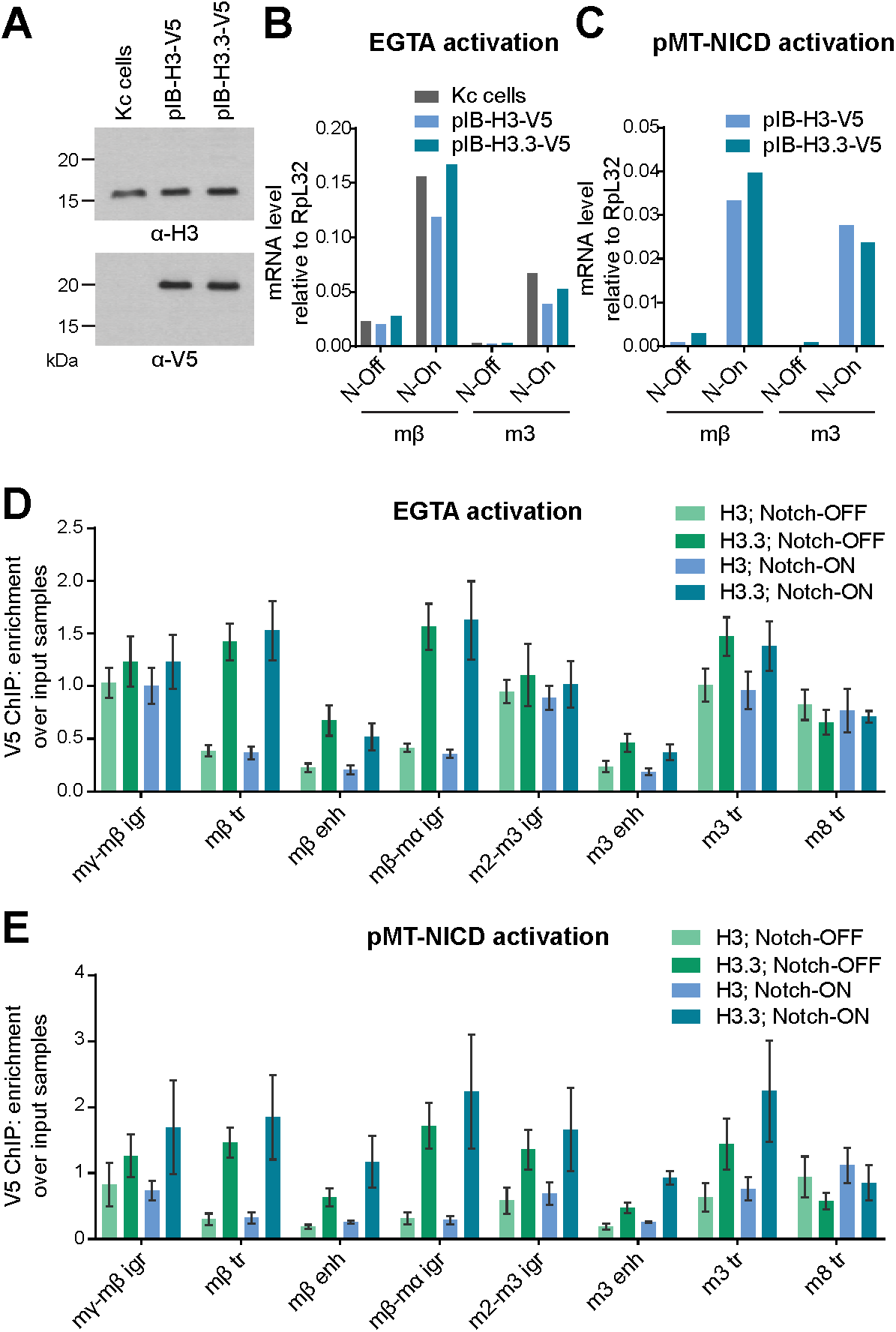
Notch activation does not affect the distribution of histones H3 and H3.3. A H3-V5 and H3.3-V5 expression in stable cell lines compared to un-transfected “Kc cells”, demonstrated by Western blots probed with H3 and V5 antibodies. V5-tagged histones have a larger molecular weight and are not detectable in the H3 blot due to low levels of expression in comparison to endogenous H3. B, C Effect of Notch activation by EGTA (B) or copper-inducible NICD expression (C) on expression of *E(spl)mβ-HLH* and *E(spl)m3-HLH* in stable cell lines expressing H3-V5 and H3.3-V5, measured by reverse transcription-qPCR. Both methods of activation strongly induce both genes. “N-On” denotes EGTA or copper treatment and “N-Off” denotes PBS alone or no copper. D, E Notch activation does not affect H3 and H3.3 levels across the *E(spl)-*C. V5 ChIP-qPCR in Kc cells expressing H3-V5 or H3.3-V5 from a ubiquitous promoter with Notch signaling activated by EGTA (D) or 6 hours of copper-inducible NICD expression (E), shown as fold enrichment over input samples. Notch activation causes no detectable change in levels compared to controls treated with PBS (D) or no copper (E). Mean +/-SEM; n = 3 (D) and 2 (E).

**Figure E5.**
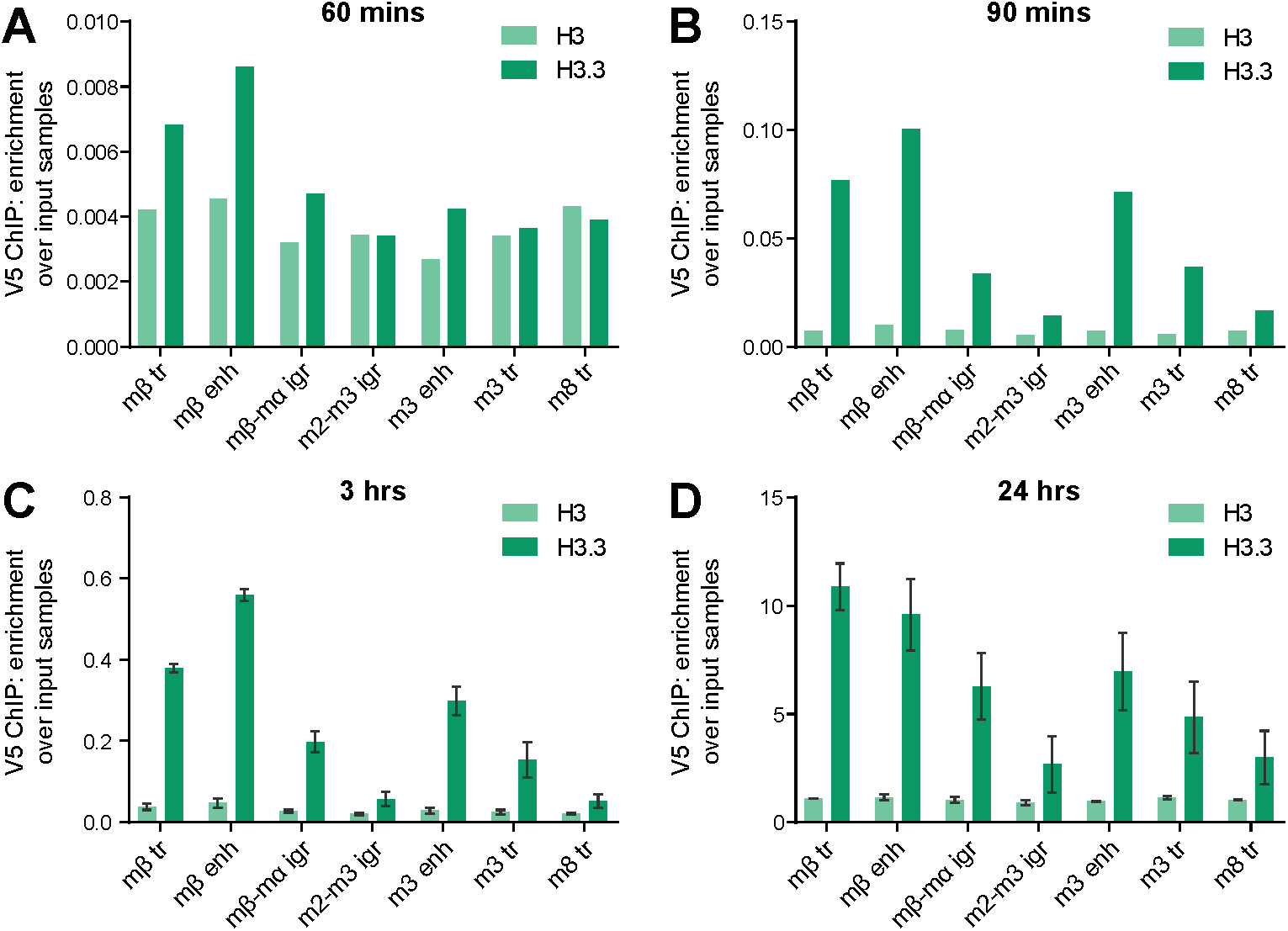
Time-course of copper-inducible histone-V5 expression. V5 ChIP-qPCR in Kc cells with H3-V5 and H3.3-V5 expression induced by copper from the pMT promoter for 60 minutes (A), 90 minutes (B), 3 hours (C) and 24 hours (D), shown as fold enrichment over input samples. Differential incorporation of H3.3 across the *E(spl)-C* is clear after 90 minutes. (C and D) Mean +/-SEM; n = 2.

